# NK Cells Engineered with a Chimeric Antigen Receptor Delay HIV Rebound and Reshape HIV Reservoir Composition

**DOI:** 10.64898/2026.01.22.700964

**Authors:** Yuan Shi, Izra Abbaali, William Harvey, Matthew Kostelny, Hongying Chen, Alexander Gonzalez, Melanie Dimapasoc, Christopher Seet, Catherine A. Blish, Jerome A. Zack, Jocelyn T. Kim

**Affiliations:** Department of Molecular and Medical Pharmacology, University of California Los Angeles, Los Angeles, CA, 90095, USA; Department of Medicine, Division of Infectious Diseases, University of California Los Angeles, Los Angeles, California, 90095, USA; Department of Microbiology, Immunology, and Molecular Genetics, University of California Los Angeles, Los Angeles, California, USA; Department of Medicine, Division of Hematology and Oncology, University of California Los Angeles, Los Angeles, California, USA; Department of Medicine, Division of Infectious Diseases and Geographic Medicine, Stanford University, Stanford, CA, USA

## Abstract

Durable HIV remission will require strategies that eliminate or permanently silence the latent reservoir that persists despite antiretroviral therapy (ART). Natural killer (NK) cells possess inherent antiviral activity, but unmodified NK cells have limited ability to recognize or clear latently infected cells during ART suppression. We engineered allogeneic human primary NK cells expressing a truncated CD4-based chimeric antigen receptor (D1D2-CAR) that targets the conserved CD4 binding site on HIV Env without permitting viral entry, and evaluated their activities in humanized mice infected with barcoded CCR5-tropic HIV. D1D2-CAR NK cells selectively killed HIV-infected primary CD4 T cells in vitro and significantly delayed viral rebound following ART interruption in humanized mice compared to GFP-NK or no NK control groups. Barcoded HIV tracking showed that CAR-NK treatment reduced the number, diversity, and inter-organ overlap of rebounding viral RNA and proviral DNA lineages, yielding rebound driven by a restricted set of dominant clones. Integration site and chromatin analysis further demonstrated selective depletion of proviruses positioned in genes, enhancers, promoters, and open chromatin. These findings show that CAR-NK cells can target rare reactivation events during ART suppression and reshape the reservoir toward a less rebound-competent, epigenetically repressive state.

## Introduction

An estimated 39.0 million people are living with HIV (PLWH) globally^1^. Despite effective antiretroviral therapy (ART), HIV persists in a long-lived latent reservoir, primarily consisting of resting CD4+ T cells that harbor integrated, replication-competent proviruses^2,3^. Viral rebound occurs rapidly upon ART interruption, and to date, the only documented cures have required allogeneic stem cell transplantation from donors homozygous for the CCR5Δ32 mutation, an approach that is neither scalable nor broadly applicable in its current format due to its risk and reliance on compatible donors^4^. Numerous alternative strategies, including latency reversing agents (LRAs), immune modulation and broadly neutralizing antibodies (bNAbs), have shown meaningful virologic impact and even post-ART viral control in select individuals^5–9^, however improved strategies to selectively eliminate reservoir cells during ART suppression are still needed.

Natural killer (NK) cells demonstrate intrinsic antiviral activity and rapid response during infection with epidemiologic links to HIV control^10–15^. NK cells recognize HIV-infected cells through a diverse array of germline-encoded activating and inhibitory receptors, enabling antigen-independent detection of HIV-infected cells^16–21^. Clinical and preclinical studies have shown that NK activation by IFNα or in combination with LRAs can reduce HIV DNA levels^22^. In addition, adoptive infusion of allogeneic haploidentical NK cells has been administered safely in PLWH^23,24^. Previously, we found that combining a protein kinase C (PKC) modulator with unmodified NK cells significantly delayed viral rebound in HIV-infected humanized mice, whereas infusion of NK cells alone during ART suppression failed to alter rebound kinetics^25^. Collectively, these findings underscore both the safety and therapeutic promise of NK cells as an off-the-shelf product, while highlighting their limited efficacy when administered without additional targeting mechanisms.

One promising strategy is to redirect NK cells toward HIV-infected cells. NK cells recognize infected or stressed cells through activating receptors and can also mediate antibody-dependent cellular cytotoxicity (ADCC), yet they can be further reprogrammed with chimeric antigen receptors (CARs) to provide antigen-specific targeting analogous to CAR-T cells. Early proof-of-concept studies using NK-92 cell lines or stem cell-derived NK cells have demonstrated HIV Env-specific recognition and inhibition in vitro^26,27^, but with limited translational relevance. Autologous CAR-T cells, including both CD4+ and CD8+ CAR T cells, are now in clinical development (NCT04648046)^28^, yet such therapies face risks of exhaustion, toxicity, and viral escape. Importantly, these same limitations, together with the added risks of graft-versus-host disease and host rejection, pose substantial challenges for developing an allogeneic T-cell product for HIV. In contrast, NK cells do not support productive HIV replication, retain innate cytotoxicity as a fallback mechanism, and can be safely administered across HLA barriers without graft-versus-host disease and less risk of cytokine release syndrome. These attributes make allogeneic CAR-NK cells a promising “off-the-shelf” immunotherapeutic platform.

Here we describe allogeneic NK cells engineered with a first-generation, proof-of-concept truncated CD4 CAR (D1D2 extracellular domains), which is known to target the conserved CD4 binding site on Env while preventing viral entry or autoinfection^29^. We evaluate their ability to eliminate infected cells during ART suppression in humanized mice infected with barcoded R5-tropic HIV. We find that D1D2-CAR NK cells can eliminate reactivating HIV-infected cells during ART suppression and reshape the reservoir landscape in vivo, providing a foundation for developing NK cell-based strategies to achieve sustained viral remission without ART.

## Results

### D1D2-CAR is efficiently expressed on NK cells and enhances clearance of HIV infection in vitro

To generate CAR-NK cells specifically targeting HIV infected cells, we first expanded and activated primary NK cells by coculturing them with a K562 cell line expressing membrane bound IL-21 and 4-1BBL^30^; then, these expanded primary NK cells were transfected with mRNA encoding D1D2-CAR-GFP or GFP only as a control. We found NK cells efficiently expressed GFP and the D1D2-CAR 24 hours post-transfection, measured by flow cytometry (Figs. 1a, b). To test the anti-HIV function of these D1D2-CAR NK cells, we cocultured them with HIV NL4-3 (a CXCR4-tropic strain) infected primary autologous CD4+ T cells at a 1:1 or 0.25:1 effector-to-target ratio for 24 hours, and then assessed the frequency of remaining infected p24+ target cells by flow cytometry (Figs. 1c, d). D1D2-CAR NK cells significantly increased viral suppression compared to GFP-NK cells or control NK cells (Fig. 1e). Viral suppression was defined as the percent reduction in p24+ CD4+ T cells in NK co-cultures relative to matched cultures without NK cells, such that higher values indicate greater NK-mediated clearance. We did not detect a significant frequency of p24+ NK cells when cocultured with infected CD4+ T cells (Figs. 1f, g), consistent with the fact that the D1D2-CAR does not allow HIV autoinfection due to the truncation of the extracellular domains required for viral entry and fusion^31^. We also did not find that D1D2-CAR NK cells significantly decreased the frequency of uninfected CD4+ T cells, suggesting the D1D2-CAR NK cells specifically targeted and killed HIV infected target cells (Fig. 1h).

**Figure 1.**
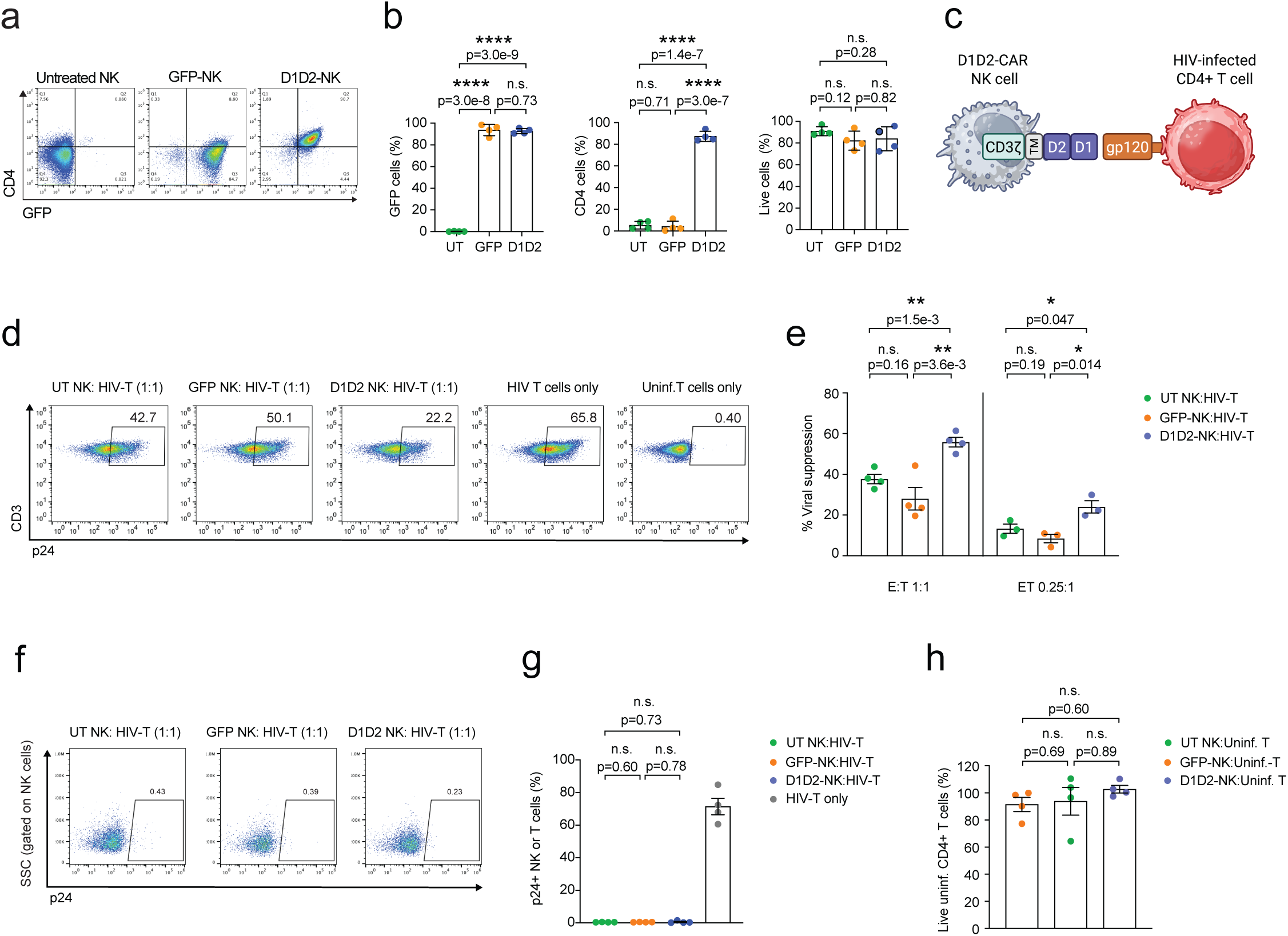
D1D2 CAR is efficiently expressed on NK cells and enhances HIV clearance in vitro. **a**) Representative flow cytometry plots of CD4 and GFP expression on expanded NK cells that were electroporated with D1D2 CAR or control GFP mRNA or no mRNA (untreated or UT) NK at 24 h post-electroporation. **b**) Percent of GFP+, CD4+ and live NK cells 24 h post-electroporation by flow cytometry. **c**) Schematic of a D1D2-CAR expressing NK cell targeting an HIV-infected cell. **d**-**g**) UT, GFP or D1D2-CAR NK were co-cultured with autologous NL4-3 infected and uninfected primary CD4+ T cells for 24 h at 1:1 or 0.25:1 E:T ratio. Representative flow cytometry plots showing frequency of p24+ T cells from cocultures containing HIV-infected T cells with UT NK cells, GFP-NK cells, or D1D2-CAR NK cells at 1:1 E:T ratio or from cultures with only HIV-infected T cells or uninfected T cells (**d**). Viral suppression or percent reduction in p24+ CD4+ T cells in NK co-cultures relative to matched cultures without NK cells (**e**). Representative flow cytometry plots showing frequency of p24+ NK cells after coculture with HIV-infected T cells at 1:1 E:T at 24 h (**f**). Percent of p24+ NK cells after 24 h of coculture with HIV-infected T cells, shown alongside the percent of p24+ T cells in cultures without NK cells (**g**). **h**) Percent survival of uninfected CD4+ T cells was calculated by comparing the frequency of live uninfected T cells in cocultures with UT NK cells, GFP-NK cells, or D1D2-CAR NK cells to the frequency of live uninfected T cells in matched cultures without NK cells. n represents 3-4 independent biological donors. Shown are mean ± SEM. P-values were determined by unpaired t test (**b**, **e**, **g**, **h**).

### D1D2-CAR NK cells efficiently engraft in vivo

While NK cells are already known to exhibit antiviral activity during productive infection and viral spread, it remains unclear whether engineered NK cells can effectively target infected cells during ART-mediated viral suppression. In this study, we specifically evaluated whether D1D2-CAR NK cells could enhance in vivo clearance of HIV-infected cells during ART suppression and early analytic treatment interruption (ATI). Our previous work demonstrated that unmodified NK cells administered after ATI can control viral rebound, but when infused during ART suppression or on the day of ATI, they did not significantly delay time to viral rebound^25^. Therefore, we asked whether D1D2-CAR expression could improve the ability of NK cells to recognize and eliminate infected cells during ART suppression or early ATI reactivation.

To this end, humanized mice were generated by intra-hepatically transplanting human CD34+ hematopoietic stem cells (HSCs) into immunodeficient NSG pups, which have minimal endogenous NK cell levels^32^. Approximately 10 weeks post-CD34+ HSC injection, human immune cell engraftment was achieved. Mice were then injected with barcoded CCR5-tropic NFNSX, which contains a genetic 21 nt tag inserted between env and nef^33^. To quantify plasma viral loads, animals were bled every two weeks post-HIV injection to confirm primary infection by RT-PCR. After 4 weeks of primary infection, animals were administered ART via oral feed for 6 weeks, which consists of raltegravir, emtricitabine, and tenofovir disoproxil fumarate (Fig. 2a).

**Figure 2.**
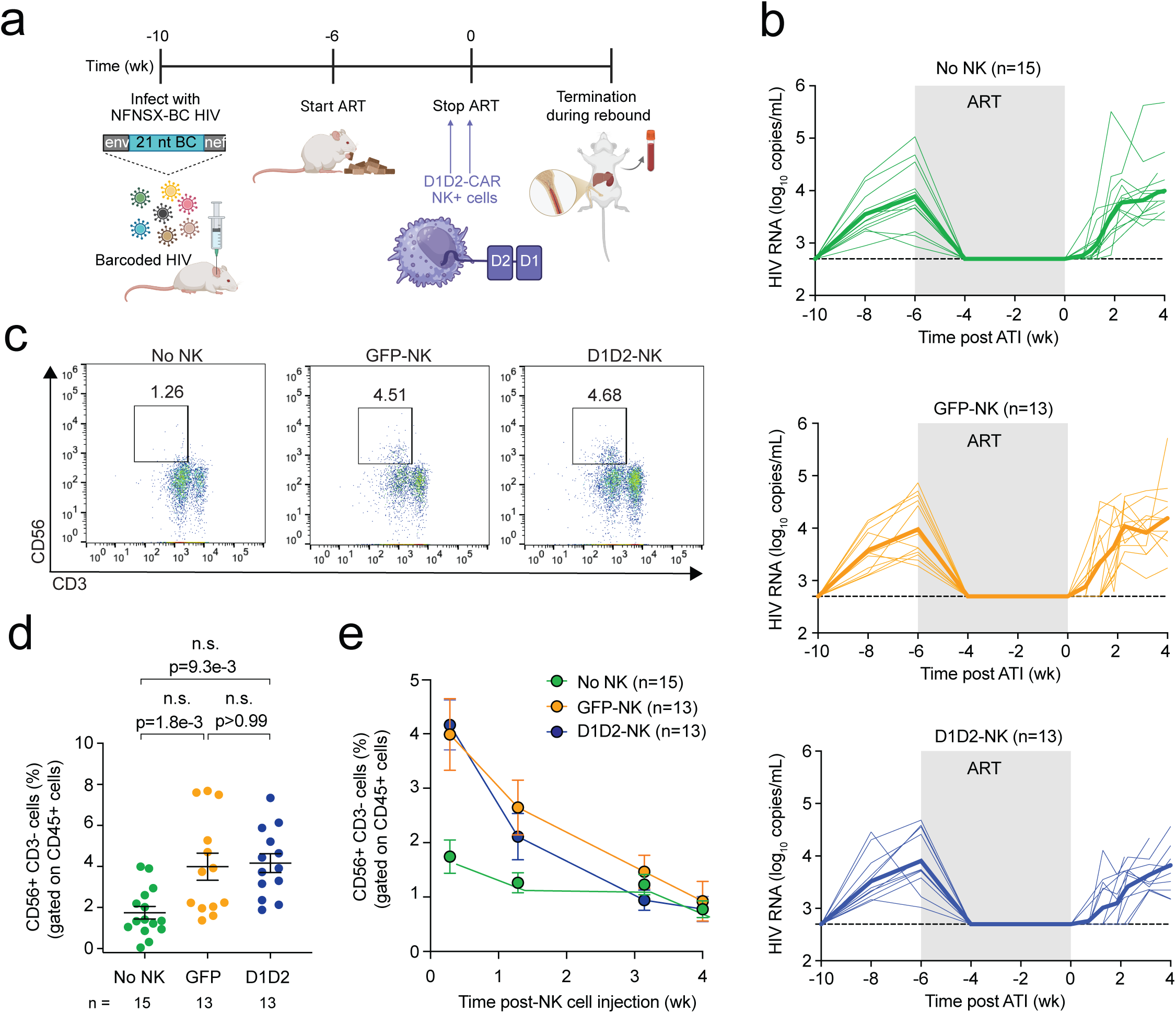
In vivo engraftment, kinetics, and tissue distribution of infused NK cells. **a**) Schematic showing CD34-NSG humanized mice infected with barcoded R5-tropic NFNSX for 4 weeks, then placed on ART for 6 weeks. 5 x 10^6^ NK cells were intravenously injected 1 week before and on the day of analytic treatment interruption (ATI). **b**) Longitudinal plasma viral loads for each infected animal at various timepoints in groups of animals that received no NK cells (green), GFP-NK cells (orange), or D1D2-CAR NK cells (blue). Thick lines represent the median viral load. Gray shading indicates an ART treatment period of 6 weeks. Black dotted line indicates the detection limit of 2.3 log RNA copies/mL. **c**) Representative Flow cytometry plots showing the frequency of human CD56+ CD3-NK cells gated among human CD45+ cells from peripheral blood of mice 2 days after the second NK cell or PBS injection. **d**) Frequency of peripheral human CD56+ CD3- NK cells gated among treated mice 2 days after the second NK cell or PBS injection. **e**) Longitudinal frequency of human CD56+ CD3- NK cells gated among human CD45+ cells from the peripheral blood of mice. n represents the number of mice (**b**, **d**, **e**). Shown are mean ± SEM (**d**, **e**). P-values were determined by two-tailed Mann-Whitney test (**d**).

After 5 weeks of ART, mice received an intravenous infusion of five million D1D2-CAR NK cells or GFP-NK cells. Control animals that did not receive NK cells received PBS vehicle injections. A second dose of NK cells was administered one week later, immediately prior to ATI. After ATI, animals were bled every week to assess time to rebound based on viral loads (Fig. 2b). Animals were sacrificed between 2 to 4 weeks after ATI once viral loads confirmed rebound infection.

We next assessed the level of human NK cell engraftment in vivo. Two days after the final dose of NK cells and ATI, the frequency of CD56+ CD3-cells NK cells were measured in the blood. As expected, we found a significantly higher frequency of NK cells among animals that received D1D2-CAR or GFP-NK cells compared to animals that did not receive NK cells (Figs. 2c, d). However, NK cell frequencies in both GFP-NK and D1D2-CAR groups declined to background levels in the blood within one to three weeks after the final injection (Fig. 2e). We then evaluated whether NK cell treatment affected viral rebound dynamics.

### D1D2-CAR NK cells delay viral rebound after ART interruption in vivo

Consistent with our hypothesis, animals that received D1D2-CAR NK cells exhibited a significant delay in time to viral rebound compared to both GFP-NK and control no NK groups (Fig. 3a). This demonstrates that engineered CAR-NK cells can act directly on either latently infected cells during ART suppression or early-reactivating cells during early ATI. Importantly, this delay was achieved with only two infusions of CAR-NK cells one week prior to and on the day of ATI, highlighting the promise of this approach in a stringent humanized mouse model where viral rebound is typically rapid and uniform. By contrast, animals receiving GFP-NK cells or no NK cells rebounded at approximately two weeks post-ATI. As expected, the time to rebound in the GFP-NK group did not differ from the no NK control group, consistent with our prior finding that NK cells without additional engineering for HIV targeting, when administered during ART suppression have minimal to modest impact on rebound kinetics^25^. At time of necropsy, the plasma HIV RNA levels (Fig. 3b), as well as the cell-associated viral RNA or total HIV DNA levels in the spleen, bone marrow, and liver (Figs. S1a, b) were similar among all groups, suggesting that once rebound occurred, viral replication was efficient. We found no significant difference in pre-ART HIV RNA levels between three groups of mice (Fig. 3c) or baseline human immune cell engraftment (Figs. 3d-f), suggesting differences in pre-ART viremia or human immune engraftment were not likely contributing factors for delayed rebound in animals receiving D1D2-CAR NK cells. NK cell engraftment in the organs was not significantly different among the three groups during rebound infection at time of necropsy (Fig. 3g), suggesting differences in NK cell engraftment likely did not account for the inferior performance of GFP-NK cells relative to D1D2-CAR NK cells.

**Figure 3.**
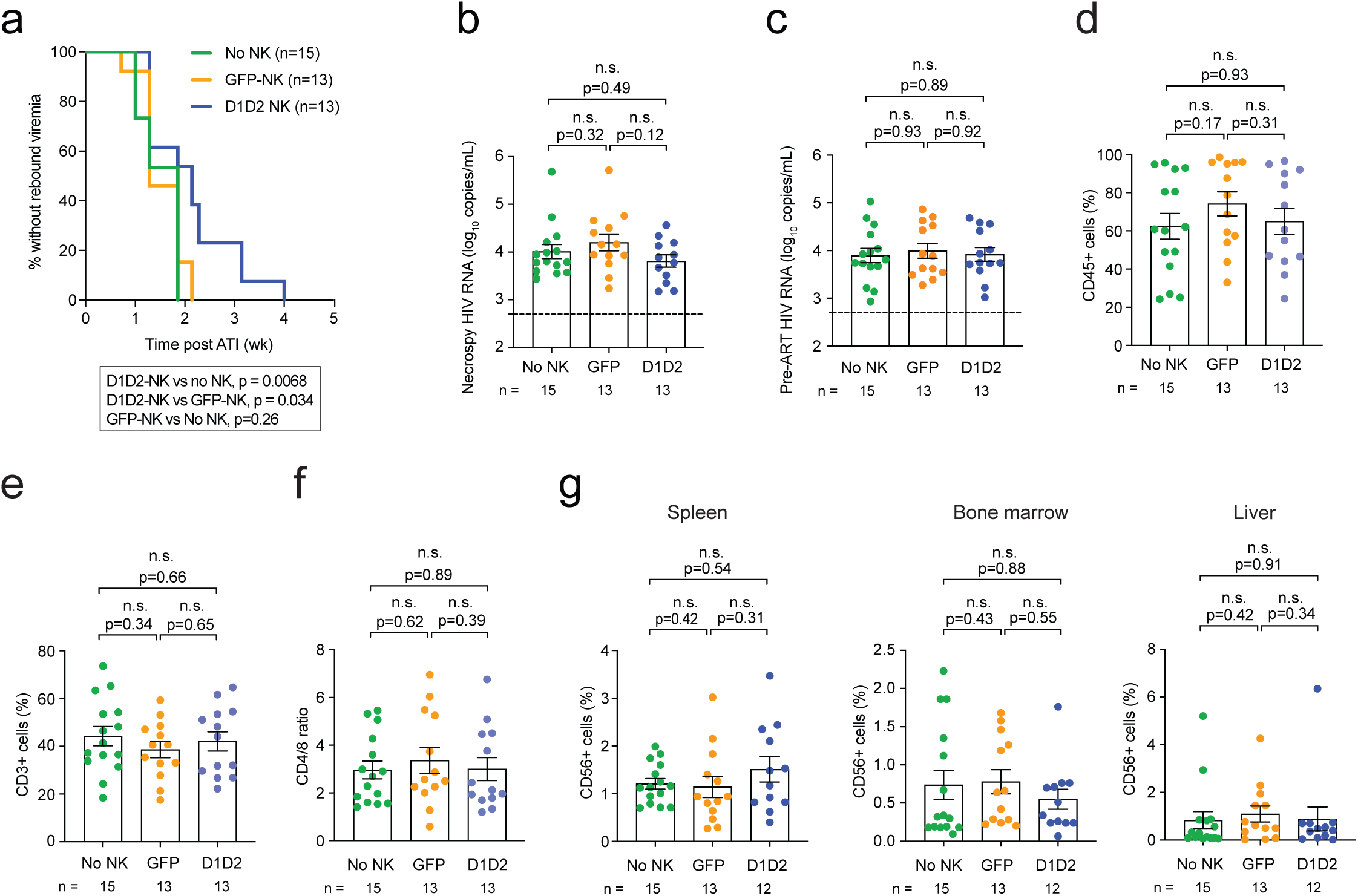
D1D2 CAR-NK delays viral rebound after ART interruption in vivo. **a**) Kaplan Meier curves of time to rebound. **b**, **c**) Plasma viral loads during rebound infection at necropsy (**b**) and before ART was initiated (**c**). **d**-**f**) Human immune engraftment was assessed by measuring the frequency of CD45+ cells (**d**), CD3+ cells (gated on CD45+) (**e**), and CD4/CD8 ratio (gated on CD3+ T cells) prior to HIV injection (**f**). Black dotted line indicates the detection limit of 2.3 log RNA copies/mL (**b**, **c**). n represents the number of mice (**a-g**). Shown are mean ± SEM (**b**-**g**). P-values were determined by log-rank (Mantel-Cox) (**a**) and two-tailed Mann-Whitney test (**b**-**g**).

### D1D2-CAR NK cells reduce the number of rebounding viral RNA barcodes

Next, we examined whether the treatment-induced delay in rebound was accompanied by a reduction in the diversity of rebounding viral lineage. Our previous studies demonstrated that a smaller reservoir is associated with delayed time to rebound following ATI, which correlates with a reduced number of rebounding viral RNA lineages^25,34^. Based on this, we hypothesized that animals receiving D1D2-CAR NK cells would reduce the number and diversity of rebounding viral RNA lineages. Here, we incorporated unique molecular identifiers (UMIs) during reverse transcription to label individual RNA molecules, amplified the viral RNA barcode region by PCR, and performed deep sequencing for analysis to assess the number and frequency of viral RNA lineages. We detected a total of 3,474 unique viral RNA barcodes from 157 samples. Consistent with the delay in viral rebound, we found D1D2-CAR NK cells significantly decreased the total number of rebounding viral RNA barcodes (Figs. 4a, b) as well as across each tissue compartment including the spleen, bone marrow, and liver compared to GFP-NK cells or no NK cells (Figs. 4c, S2a-c). As anticipated, control GFP-NK cells did not significantly decrease the number of rebounding viral clones compared to animals that did not receive NK cells.

**Figure 4.**
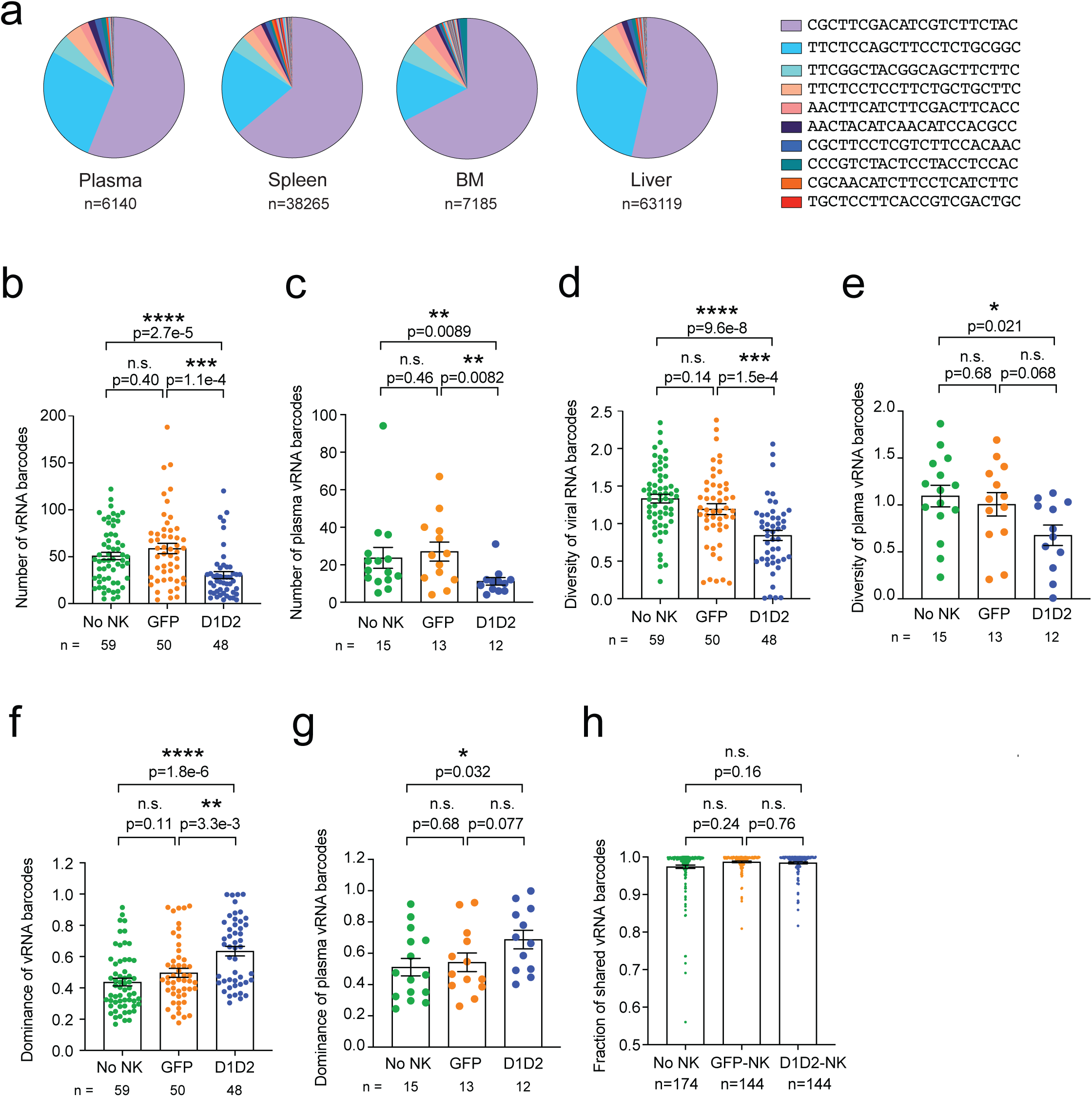
D1D2-CAR NK cells reduce the number and diversity of rebounding viral barcodes. **a**) Representative pie charts depicting distribution of viral RNA barcodes in the plasma, spleen, BM and liver of one mouse. Genetic sequence of ten barcodes shared among the organs is shown in the key. **b**, **c**) Number of rebounding viral RNA barcodes in all organs (**a**) and in the plasma (**b**) quantified by deep sequencing by treatment group. **d**, **e**) Genetic diversity rebounding viral RNA barcodes from all organs (**d**) and plasma (**e**) measured using Shannon’s diversity. **f**, **g**) Dominance of viral RNA barcodes measured using Simpson’s index. n represents the number of organs. **h**) Fraction of shared viral RNA barcodes between organ pairs within an animal was quantified by overlap index. n represents the number of organs (**b**, **d**, **f**), mice (**c**, **e**, **g**), and organ pairs (**h**). Shown are mean ± SEM. P-values by two-tailed Mann-Whitney test.

### D1D2-CAR NK cells reduce diversity and increases dominance of viral RNA barcodes

We quantified the genetic diversity of rebounding viral barcodes using Shannon’s entropy, which captures both the richness and evenness of contributing lineages. A high entropy reflects a diverse population with many lineages contributing to rebound, whereas a low entropy indicates a more clonal rebound dominated by fewer lineages. We found mice treated with D1D2-CAR NK cells exhibited significantly lower diversity of the viral barcodes compared to mice that received GFP-NK cells or no NK cells (Figs. 4d, e, S2d-f), suggesting that D1D2-CAR NK cells depleted not only the total number of viral barcodes, but also their genetic richness. To further characterize the composition of the rebound population, we measured Simpson’s dominance, which quantifies the extent to which one or a few barcodes dominate the viral population. Animals treated with D1D2-CAR NK cells displayed significantly higher dominance indices than controls (Figs. 4f, g, S2g-i), demonstrating that rebound in these animals was driven by a small set of viral clones. Together, these findings show that D1D2-CAR NK cells reduce overall barcode count and diversity, leaving rebound to emerge from a restricted set of dominant viral lineages.

### D1D2-CAR NK cells do not limit dissemination of viral RNA barcodes following rebound

We next examined whether rebounding viral barcodes disseminated evenly across the organs including plasma, spleen, bone marrow, and liver. To do this, we calculated an abundance-weighted fractional overlap index (referred to as overlap index in the following), which quantifies the proportion of barcodes shared between any two organs within the same animal. A value of 1.0 indicates complete overlap, whereas 0 indicates no overlap. We found the fraction of shared viral RNA barcodes was similar across all groups and broadly distributed among organs (Fig. 4h). These data suggest that although D1D2-CAR NK cells significantly impact the reactivation kinetics of the viral reservoir, they do not limit the dissemination of viral lineages once rebound occurs, likely reflecting spread by cell-free virions.

### D1D2-CAR NK cells restrict proviral lineage number and inter-organ dissemination

Having established that D1D2-CAR NK cells restrict the number, diversity, and dominance of rebounding viral RNA lineages, we next asked whether these effects were reflected at the level of the proviral reservoir. Here we quantified each HIV DNA molecule by UMI and used PCR linkage to simultaneously amplify the proviral barcode and integration site^33^. We then focused on proviral barcode analysis and examined both the total number of proviral barcodes and the subset of proviral barcodes that had a matching barcode detected in plasma viral RNA during rebound. As in our prior study, we classified a provirus as ‘associated with viremia’ when its barcode was also detected in plasma viral RNA at rebound, representing clones that both reactivated and successfully reseeded tissues. Out of a total of 908 unique proviral barcodes, we detected 88 proviral barcodes associated with viremia. Similar to what we observed for viral RNA barcodes, the number and diversity of proviruses associated with viremia or total proviruses were significantly lower in animals receiving D1D2-CAR NK cells compared to animals receiving GFP-NK cells or no NK cells (Figs. 5a, b, S3a, b). In addition, the dominance of proviral barcodes was higher in the D1D2-CAR NK cell-treated group compared to the GFP-NK group and no NK control group (Figs. 5c, S3c). These findings suggest that D1D2-CAR NK cells restrict the number of viral lineages associated with rebound viremia at the proviral level as well.

**Figure 5.**
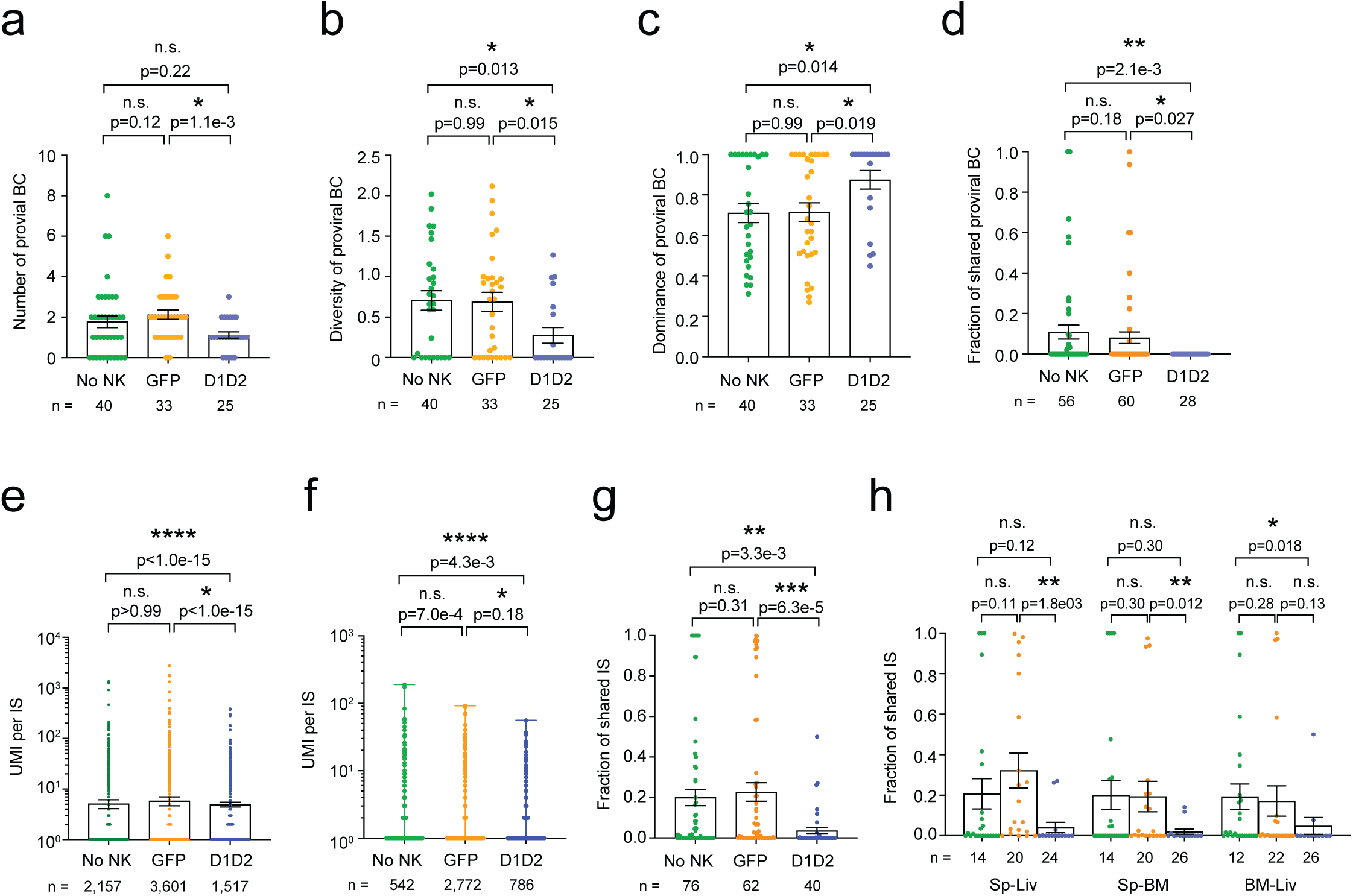
D1D2-CAR NK cells reduce proviral viral barcodes associated with rebound infection and sharing of integration sites. a-c) The number (**a**), genetic diversity (**b**) and dominance (**c**) of proviral barcodes associated with rebound viremia were quantified by deep sequencing. **d**) The fraction of shared proviral barcodes associated with rebound viremia between organ pairs were quantified by overlap index. **e**, **f**) The size of expanded cell clones harboring any provirus (**e**) or those with a provirus associated with rebound viremia (**f**) was quantified by the ratio of UMI per integration site (UMI/IS) using deep sequencing. **g**, **h**) The fraction of shared integration sites by treatment group (**g**) and by organ pairs (**h**) was quantified by overlap index. n represents the number of organs (**a**-**c**), integration sites (**e**, **f**), and organ pairs (**d**, **g**, **h**). Shown are mean ± SEM. P-values by two-tailed Mann-Whitney test.

### D1D2-CAR NK cells limit inter-organ dissemination and intra-organ expansion of infected cell clones

We next examined whether D1D2-CAR NK cells altered the tissue dissemination patterns of these rebound-competent clones by quantifying the frequency of shared proviral barcodes across different organs within the same animal using the overlapping index. We found the fraction of shared proviral barcodes associated with viremia and total proviral barcodes was significantly lower in animals receiving D1D2-CAR NK cells compared to animals receiving no NK cells, and trended lower relative to GFP-NK-treated animals (Figs. 5d, S3d). A trend towards reduction in shared proviral barcodes for each organ pair was observed among animals receiving D1D2-CAR NK cells compared to GFP-NK or no NK groups (Fig. S3e). Collectively, these results suggest that D1D2-CAR NK cells not only reduce the number of rebound-competent clones but also limit their intra-organ dissemination.

We have previously described how the ratio of the UMI for each integration site (UMI per IS) reflects infected cell clone size because each UMI corresponds to a distinct integration site molecule derived from a distinct infected cell^33^. To assess whether D1D2-CAR NK cells could affect the size of infected cell clones, we analyzed the UMI per IS ratio, which is a readout of cell clone size. When we looked at all integration sites or integration sites linked to a proviral barcode associated with rebound viremia, we found that UMI per IS was significantly lower in animals that received D1D2-CAR NK cells compared to GFP-NK or no NK controls (Figs. 5e, f). Interestingly, this difference was even more pronounced when we examined the subset of integration sites with UMI per IS > 1, representing cell clones that were detected as larger than a single cell (Fig. 4Sa). This pattern also held when we looked at the expanded clones linked to proviral barcodes associated with rebound viremia (Fig. S4b).

We then assessed the distribution of these clones by measuring the fraction of shared integration sites between pairs of organs within the same animal, using the overlap index as a measure of clonal overlap. Notably, the fraction of shared integration sites was significantly lower in animals treated with D1D2-CAR NK cells compared to those treated with GFP-NK or no NK cells (Figs. 5g, h, S4c), indicating reduced dissemination of individual cellular clones across tissue compartments.

### D1D2-CAR NK cells reshape the chromatin landscape of proviruses associated with rebound viremia

To determine if NK cell therapy reshapes the chromatin environments of rebound-competent proviruses, we focused on proviruses that were detected in plasma viral RNA during rebound viremia. Because HIV preferentially integrates within active transcription units^35,36^, we first aligned integration sites to the human genome (hg38, Ensemble release 108) from the USCSC Genome Browser database^37^ and assessed whether proviruses differed in their genetic distribution across treatment groups and then extended this analysis to a broader definition of transcriptionally active regions, defined here as genes, promoters, enhancers, and DNase-accessible open chromatin. Mosaic plots showed that the odds of a provirus being located within a gene was 0.72-fold (28% lower odds) in D1D2-NK treated animals compared to no NK controls, whereas GFP-NK treatment produced no difference (Fig. 6a). Using the broader active chromatin definition, proviruses in D1D2-treated animals were also significantly less likely to be in active chromatin compared to no NK controls (odds ratio = 0.68; 32% lower odds), with again no detectable change in GFP-NK animals relative to no NK controls (Fig. 6b). These results suggest a pattern of selective pressure specific to D1D2-CAR NK cells, which eliminates proviruses integrated in genic or otherwise active regions, leaving behind a reservoir enriched for proviruses in non-genic or inactive regions.

**Figure 6.**
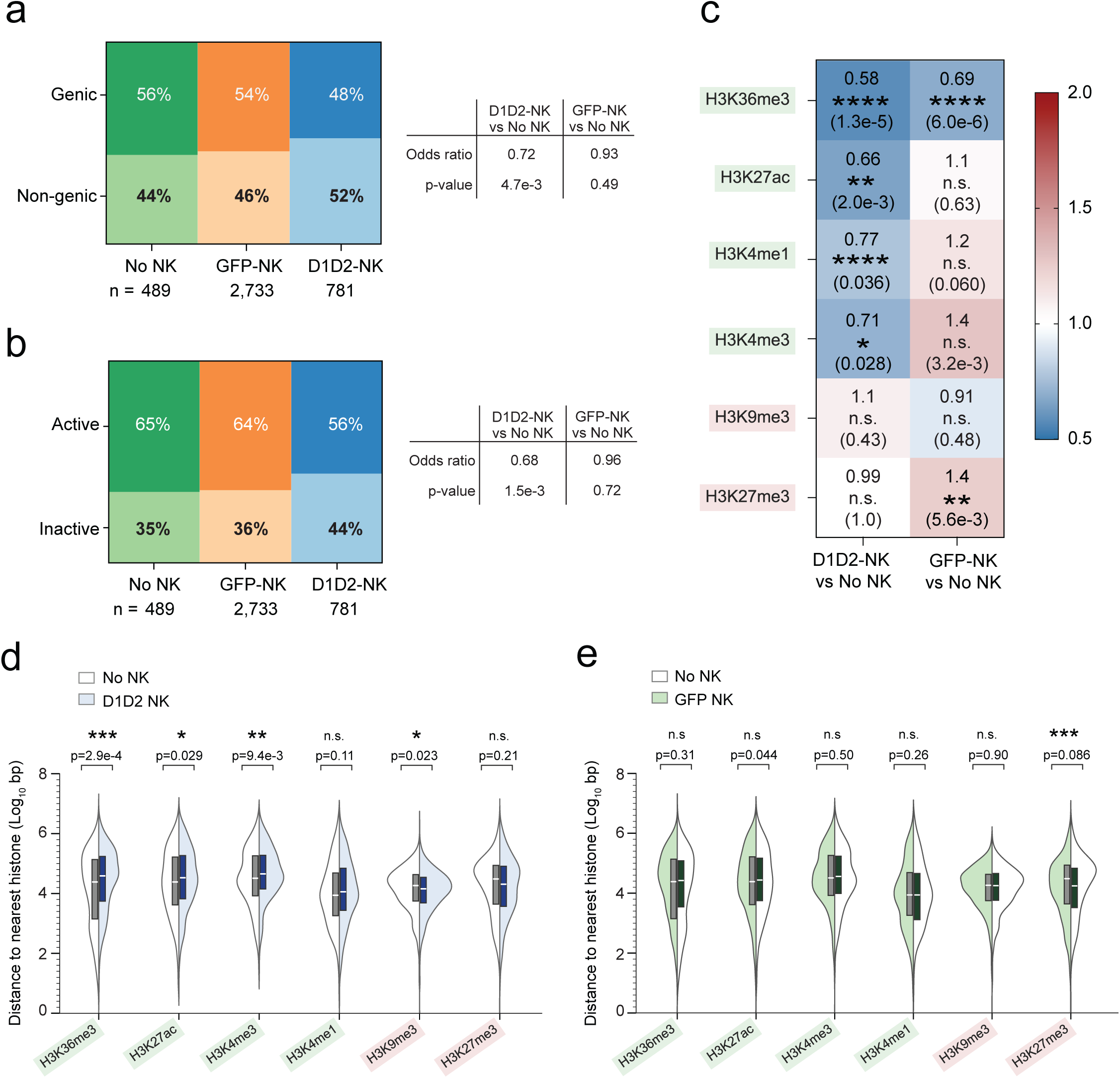
D1D2 CAR-NK cells alter chromatin contexts of rebound-associated proviruses. **a**, **b**) Mosaic plots showing the distribution of proviruses associated with rebound viremia integrated within genic versus non-genic regions (**a**) or within transcriptionally active (genes, promoters, enhancers, or DNase-accessible regions) versus inactive regions (**b**) across treatment groups. Odds ratios and p-values determined by two-tailed Fisher exact test. **c**) Heatmap showing fold-change ratios and significance values for distribution of rebound-associated proviruses in activating (H3K36me3, H3K27ac, H3K4me1, H3K4me3) or repressive (H3K9me3, H3K27me3) histone regions, based on ChIP-seq annotations from activated primary CD4 T cells in ENCODE. Values reflect the relative likelihood of overlap with each histone mark for each NK treatment arm compared to No NK controls. (**d**, **e**) Violin plots comparing the distance of rebound-associated proviruses to the nearest activating or repressive histone mark for D1D2-CAR NK vs No NK (**d**) or GFP-NK vs No NK (**e**) groups. Shown are mean ± SEM. P-values were calculated by two-tailed Mann-Whitney test.

We next assessed whether histone modifications surrounding proviruses associated with rebound viremia differed across treatment groups. We aligned the integration sites to the annotated chromatin immunoprecipitation sequencing (CHIP-seq) deep sequencing data of activated primary CD4 T cells histone modifications in the Encyclopedia of DNA element (ENCODE) regions^38^ and compared the relative proximity of each rebound-associated provirus to major activating and repressive histone modifications. Activating marks included H3K36me3, a transcriptional elongation mark enriched across gene bodies, and the enhancer- and promoter-associated marks H3K27ac, H3K4me1 and H3K4me3. Repressive marks analyzed were H3K9me3, associated with heterochromatin-mediated HIV silencing^39^,and H3K27me3, associated with Polycomb-mediated repression^40,41^. We calculated fold-change ratios in the distance to each mark between treatment groups and assessed their significance using two-sided Fisher tests.

We then quantified whether proviruses in the D1D2-NK group were less likely to be located within regions marked by activating histone modifications (H3K36me3, H3K27ac, H3K4me1 and H3K4me3), and, for those proviruses outside these regions, whether they resided farther from these histone marks compared to proviruses in No NK controls. We found that proviruses associated with rebound viremia in D1D2-treated animals were significantly less likely to overlap activating histone marks (Fig. 6c) and were positioned farther away compared to proviruses in the No NK group (Fig. 6d). In contrast, GFP-NK treatment produced no significant differences for any activating mark except H3K36me3 (Fig. 6c), though without a significant difference in distance compared to No NK (Fig. 6e). For the repressive histone marks, the D1D2-CAR NK arm showed a modest increase in proximity to H3K9me3 (Fig. 6d), while GFP-NK proviruses were more likely to overlap or reside closer to H3K27me3 (Figs. 6c and 6e), suggesting that both NK treatments may exert mild selective effects on proviruses positioned near repressive chromatin, though not with the magnitude or consistency observed for activating marks. Collectively, these findings suggest that D1D2-CAR NK cells selectively eliminate proviruses positioned in regions associated with transcriptional activation, leaving behind a reservoir enriched in proviruses located within more inaccessible or repressive chromatin contexts.

## Discussion

Natural killer (NK) cells have long been implicated in HIV control^17,42–46^, however their function is often impaired and not completely restored by ART^42,47,48^. Infusion of unmodified NK cells has shown limited to modest ability to recognize or clear latently infected cells during ART, when antigen expression is minimal^24,25,49^. Here we demonstrate that D1D2-CAR NK cells can significantly delay HIV viral rebound following ART interruption and reshape the composition of the viral reservoir in vivo. By combining CAR engineering with the innate antiviral properties of NK cells, we show that a single antigen-specific D1D2 CAR design redirects NK cell cytotoxicity toward HIV-infected targets without triggering nonspecific depletion of uninfected CD4+ T cells or autoinfection of the effector cells themselves. These findings suggest that D1D2-CAR NK cells may overcome several limitations associated with current HIV cure strategies, including poor specificity and ineffective clearance of latently infected cells during ART suppression.

An important distinction of this study is that CAR-NK efficacy was evaluated under ART suppression where unmodified NK cells are known to have minimal or modest impact. We have previously shown that NK infusions administered after ATI can transiently control viral rebound, whereas NK cells given during ART suppression do not significantly alter rebound timing^25^. The ability of D1D2-CAR NK cells administered during ART to delay post-ATI rebound therefore contrasts sharply with the lack of benefit observed with non-CAR NK cells in this and prior studies. Although unmodified NK cells can control actively replicating HIV, their activity during ART suppression is likely limited because spontaneously reactivating proviruses are rare and antigen availability is low. In our model, the first NK infusion is given during full ART suppression, when antigen is likely low or transient, and the second infusion occurs on the day ART is stopped, when only the earliest, sporadic reactivation events are beginning. Under these conditions, D1D2-CAR NK cells must identify and eliminate rare, newly reactivating infected cells before they reseed the reservoir, representing an exceptionally stringent functional test.

The comparable pre-ART viral loads, baseline immune reconstitution, and NK persistence across groups indicate that the therapeutic benefit likely arises from the Env-specific D1D2-CAR recognition rather than differences in viral replication, humanization of the mice, or engraftment of adoptively transferred NK cells. Collectively, these findings highlight CAR engineering as the crucial modification that enables NK cells to engage the reservoir during ART suppression and eliminate infected cells in a window where unmodified NK cells demonstrate limited efficacy.

The reservoir analyses provide mechanistic insight into how CAR-NK cells reshape rebound viremia. Using a barcoded HIV system, we found that D1D2-CAR NK cells markedly reduced the number of rebounding viral RNA barcodes, decreased clonal diversity, and increased dominance, indicating that rebound originated from a reduced set of viral lineages. Importantly, CAR-NK treatment also reduced proviral barcode number and decreased inter-organ dissemination of infected cell clones, suggesting that CAR-NK cells not only eliminate reactivating proviruses but also restrict the tissue spread of infected cells. These findings are consistent with recent work showing that haploidentical NK cell infusion in adults reduced the frequency of HIV RNA positive cells in lymph node and gut tissues^24^, supporting the concept that NK cell-based interventions can directly constrain tissue reservoirs. This pattern resembles recent human data showing that NK cell related immune programs can reduce the viral reservoir in individuals who acquired HIV perinatally and initiated ART very early^50^, supporting the idea that a functional NK response can help shape the reservoir during ART or early rebound.

A mechanistic hallmark of this study is the discovery that D1D2-CAR NK cells impose selective pressure based on the proviral chromatin context. Proviruses associated with rebound viremia in CAR-NK treated animals were significantly less likely to occupy genic regions, promoters, enhancers, or open chromatin, and were less likely to align in and around activating histone marks such as H3K27ac, H3K4me1/3, and H3K36me3^51,52^. This contrasts with well-established features of HIV biology; proviruses preferentially integrate into active chromatin^35,36,53^ and clonally expanded or transcriptionally active proviruses are frequently located near genes and epigenetic features associated with active chromatins^54–56^. These findings suggest that CAR-NK therapy disproportionately eliminates infected cells harboring proviruses in accessible, transcriptionally active chromatin, which could be the cells most likely to reactivate early during ART release. Conversely, the proviruses that persist after CAR-NK treatment are more likely to be associated with inaccessible or repressive chromatin contexts, resembling a “deep latency” state^57^. This selective survival pattern provides a conceptual framework for how antigen-directed NK responses could remodel the epigenetic architecture of the reservoir and potentially synergize with future latency-reversing interventions^58^.

These results also complement and extend emerging clinical studies evaluating NK cell therapies in PLWH. Multiple clinical trials have demonstrated that allogeneic NK cell infusion is safe^59–61^ including for PLWH^23,24^. NK activation by IFN-α or latency reversing agents (LRAs) enhance HIV-infected cell clearance^25^, and bNAbs rely heavily on NK-mediated ADCC for therapeutic efficacy^62^. Meanwhile, CAR-NK cells have shown excellent safety and promising therapeutic efficacy in oncology^59,63,64^, however prior pre-clinical work using iPSC-derived NK cells expressing a CD4-ζ CAR demonstrated enhanced in vitro recognition of HIV-infected targets, but no added suppression of HIV replication during acute infection of humanized mice^26^. In contrast, the present study shows that a proof-of-principle truncated CD4-ζ-based CAR expressed on NK cells provides modest but measurable benefit under stringent, low-antigen conditions such as during ART suppression or early ATI, suggesting that more optimized CAR-NK designs could outperform unmodified NK cells during this window.

Several limitations warrant consideration. First, the humanized mouse model, while highly informative, does not fully recapitulate lymphoid architecture, NK cell homeostasis, or reservoir complexity in the human body. While other humanized mouse models may support partial lymphoid architecture, it is not clear whether NK activity in these systems are overestimated as it is not expected that unmodified NK cells alone would clear the reservoir after ATI. Second, we found NK persistence was relatively short-lived, and although sufficient to alter reservoir composition, enhancing in vivo survival may improve their efficacy. Third, this study did not incorporate LRAs, checkpoint blockade, or other immunomodulators that could further sensitize the reservoir to CAR-NK clearance. Lastly, because D1D2-CAR NK cells preferentially eliminate cells harboring proviruses integrated in active or genic regions, future studies should dissect the molecular basis of this selectivity, consistent with prior work showing that transcriptionally repressed or epigenetically silenced reservoir cells can be less susceptible to CTL or NK-mediated clearance^65,66^.

In conclusion, our findings show that D1D2-CAR NK cells are a promising therapy that can selectively deplete infected cells harboring inducible HIV during ART suppression and restrict the composition of rebound viremia. By targeting early, rare reactivation events during ART, CAR-NK cells restrict rebound viremia, prune susceptible proviral lineages, limit inter-organ dissemination, and remodel the chromatin features of proviruses that persist. These attributes make D1D2-CAR NK cells an attractive candidate for combination strategies. Synergy with LRAs^25,67^, integration with bNAb therapy, or pairing with agents that enhance NK persistence may amplify reservoir clearance. Ultimately, these results advance the prospect of an off-the-shelf, antigen-specific CAR-NK approach for achieving durable ART-free remission and bring the field closer to a scalable, broadly applicable HIV cure strategy.

## Methods

### Ethics Statement

All mouse experiments were performed in compliance with the study protocol approved by the UCLA Animal Research Committee (ARC-2021-020). De-identified PBMCs from healthy human donors were obtained under informed consent from the UCLA-CDU CFAR Virology Core under IRB approval then provided to investigators in an anonymized fashion.

### Mice

All mice were maintained in the animal facility at UCLA. In brief, NOD.Cg-*Prkdcscid Il2rgtm1Wjl*/SzJ or NSG mice^32^ were bred at UCLA (RRID:IMSR_JAX:005557). Male and female pups were age-matched and received 1Gy irradiation by X-ray at day 1 to 3 of life. CD34+ human hematopoietic cells were intra-heptically injected by the UCLA humanized mouse core.

### Primary NK cells

Primary human NK cells were isolated from human PBMCs by immunomagnetic depletion via a NK cell isolation kit (Miltenyi Biotec, 130092657). NK cells were expanded by coculturing with K562 cells (ATCC, CCL-243) expressing membrane-bound human IL-21 and 4-1BBL (REF) at a ratio 1:2 for 2 weeks in C10 media (RPMI 1640 media supplemented with 10% vol/vol FBS (Omega Scientific, NC2117537), 1% L-glutamine, 1% penicillin/streptomycin (Gibco, 10378016), 500 mM 2-mercaptoethanol (Sigma, M6250), 1 mM sodium pyruvate (Gibco, 11360070), 0.1 mM MEM nonessential amino acids (Gibco, 11-140-050), 10 mM HEPES (Gibco, 15-630-080), and 20 ng/mL of recombinant human interleukin-2 (IL-2) (ThermoFisher, 200-02) and 20 ng of recombinant human interleukin-15 (ThermoFisher, 200-15).

### NK cell transfection

Primary NK cells were harvested on Day 14 of expansion and counted using the LUNA™ automated cell counter with viability dye. Cells were washed once with X-VIVO 15 media (Lonza, #02-053Q) to remove residual fetal bovine serum (FBS), as serum can reduce electroporation efficiency. Washed cells were centrifuged at 300 x g for 10 minutes, resuspended at 30 × 10⁶ cells per aliquot in microcentrifuge tubes, and incubated at 37 °C with 5% CO₂ until electroporation. To prepare the electroporation reaction, media was removed and cells were resuspended in a 1:1 mixture of cold electroporation buffer (Buffer A + B, Celetrix #1204) with 20 µg of D1D2 mRNA construct (GFP-2A-D1D2-CD3z L-7009, Trilink) in a total volume of 125 µL. The mixture was incubated on ice for 5–10 minutes prior to transfer into 125 µL electroporation cuvettes, avoiding the introduction of bubbles. The loaded electroporation tube was placed on ice for a couple of minutes followed by electroporation using the Celetrix electroporator set at 1150V, 20 ms pulse width, pulse number 1, and 1 ms pulse interval. Post-electroporation, cells were immediately transferred into 400 µl of prewarmed X-VIVO 15 supplemented with 100 ng/mL IL2 and 20 ng/mL IL15 and allowed to rest at 37 °C for at least 15 min. Cell viability was assessed using the LUNA™ cell counter. Electroporated cells were pooled, centrifuged, and resuspended in sterile PBS at 50 × 10⁶ cells/mL. For in vivo experiments, 5 × 10⁶ D1D2-electroporated NK cells were retro-orbitally injected into mice. Cells were stained for GFP+ and CD4+ expression 24 hours post-transfection to verify transfection efficiency.

### Virus production

Plasmid DNA encoding the barcoded NFNSX^33^ was transfected into HEK293T/17 cell line purchased from the American Type Culture Collection (ATCC, CRL-11268) using following the manufacturer’s protocol for the BioT transfection reagent (Bioland, B01-01). Cell-free supernatant was harvested 48 h after transfection and passed through a 0.45 µm filter. DNaseI (40ng/mL) was added to the library to remove residual DNA from the supernatant. The barcoded virus supernatant was aliquoted and frozen at −80°C for future use.

### *In vitro* co-culture experiments

PBMC were co-stimulated with Dynabead CD3/CD28 human T-activator (ThermoFisher Scientific, 11131D) per manufacturer’s instructions and cultured in C10 media supplemented with 20 ng/mL of recombinant human IL-2 for 2-3 days. CD4+ T cells are isolated from PBMCs using CD4 MicroBeads (Miltenyi, 130-097-048) according to the manufacturer’s instructions.

CD4+ are mixed with unconcentrated NL43 strain HIV-1 at an MOI of 0.16 in C10 supplemented with 20 ng/mL IL2 and 10 µg/mL polybrene. Cells are spinoculated at 1200 g for 2 hours at 25 °C with slow/no brakes. One day post-infection, 1:1 and 0.25:1 mixtures of untreated, GFP or D1D2-NK cells (effector) to infected CD4+ T cells (target) ratios are cultured to setup the functional coculture assay.

### Plasma viral load measurements

For weekly or biweekly sampling, 50 µl of blood was collected by retro-orbital bleed into EDTA-coated capillary tubes, as previously described^25,33^. Blood was centrifuged at 300×g for 5 min, and plasma RNA was extracted using the QIAamp Viral RNA Mini Kit. HIV-1 RNA was quantified by qRT-PCR using TaqMan Fast Virus 1-Step Master Mix with 20 µl of eluted RNA and gag-specific primers (HIV-1-gag_F1/R1) and probe. Viral load was determined by comparison with plasmid DNA standards, with a quantitation limit of 200 copies/mL. At necropsy, >100 µl of blood was collected to obtain ≥50 µl plasma for RNA barcode analysis.

### Blood and tissue harvest and processing

Blood was centrifuged to separate plasma and cells; plasma was used for viral load measurements, and the cellular fraction was subjected to RBC lysis to obtain PBMCs as previously described^33^. Splenocytes were isolated by mechanical dissociation through 100 µm and 40 µm filters followed by RBC lysis. Bone marrow was flushed from femurs and tibias, filtered through a 70 µm strainer, and RBC-lysed. Liver leukocytes were enriched using a 70% Percoll gradient. All cell suspensions were resuspended in R10 medium and counted using a Luna dye-based counter. Cells from blood, spleen, bone marrow, and liver were either fixed for flow cytometry, stored as pellets for DNA analysis, or resuspended in RLT buffer for RNA extraction

### Staining and analysis for flow cytometry

Cells from animals were stained with fluorescently conjugated anti-human antibodies: CD3-Pacific Blue (clone Hit3a), CD8a-FITC (clone Hit8a), CD4-PE (clone RPA-T4), CD45-APC (clone 2D1), CD56-PE-Cy7 (clone HCD56) (all from Biolegend) and Ghost Dye Red 780 (Tonbo Biosciences). All flow cytometry samples were run using an Attune NxT (Beckman Coulter). All data was analyzed using FlowJo v.10.6.0 (TreeStart, Inc) (Fig. S5).

### Cell-associated (CA)-HIV RNA and total HIV DNA

Cell-associated HIV RNA was extracted from lysed splenocytes, bone marrow cells, and human thymic implants using RNeasy Mini Kit. Genomic DNA was extracted from cell pellets using DNeasy Blood & Tissue Kits (Qiagen, 69504). Viral loads were measured by qPCR using 500 ng of CA-HIV RNA and DNA with the same primers as above. The same amount of input RNA and DNA that was used for viral load measurement was used for barcode analysis.

### RNA barcode analysis

RNA was extracted from plasma using QIAamp Viral RNA Mini Kit (Qiagen, 52904) and from cells using RNeasy Mini Kit (Qiagen, 74104). cDNA was synthesized using the SuperScript IV First-Strand Synthesis Kit with the same input RNA amount used for viral load quantification, as previously described^33^. The barcode region was amplified by hemi-nested PCR using Phusion High-Fidelity DNA Polymerase. Each RNA molecule incorporated a Primer ID during reverse transcription. Primer ID removal and cDNA purification were performed using the PureLink PCR Purification Kit. cDNA were subject to two rounds of semi-nested PCR to amplify the barcode and UMI region. Amplicons were end-repaired, A-tailed, and ligated to Illumina sequencing adapters using the NEBNext Ultra II End Repair/dA-Tailing and Ligation Modules. Ten-nucleotide multiplexing indexes on both ends were used to distinguish samples. Libraries were sequenced on an Illumina NovaSeq6000 SP (PE150) system at ≥ 10× read depth relative to viral genome copies. Reads were de-multiplexed using the sample index and filtered to remove low-quality bases (Q < 30) and mismatches between paired reads. Primer IDs were used for error correction and collapsed using a cutoff defined by the Poisson distribution of sequencing errors, as previously described. For each Primer ID, the consensus barcode was called, and barcodes differing by ≥4 bp were grouped into clusters.

### Barcode diversity analysis and abundance-Weighted Fractional Overlap Calculation

The barcode diversity was analyzed using the Shannon diversity index and Simpson dominance index as previously described^33^. To quantify the overlap of viral lineages between tissues, we calculated an abundance-weighted fractional overlap index for every pair of organs (*A* and *B*) within an animal. Shared lineages were defined as those possessing identical integration sites (IS). The overlap score was determined using the following formula:

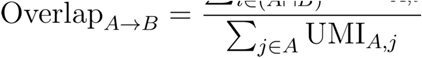

where UMI_A,i_ represents the clone size (UMI count) of a shared integration site *i* within tissue A, and the denominator represents the total proviral load (Total UMIs) in tissue A.

Overlap indices were calculated for all possible pairwise permutations of tissues within each animal.

### DNA sequencing library preparation

The barcode and integration site linkage sequencing library was prepared as previously described^33^. Briefly, DNA and RNA were isolated using the AllPrep DNA/RNA Mini Kit (Qiagen). One microgram of genomic DNA was digested with HinP1I (NEB), purified, and end-repaired with the Ultra II End Repair Module (NEB). A custom annealed adapter containing a 14-nt UMI was ligated to fragmented DNA. Adapter-ligated DNA was amplified through four rounds of semi-nested PCR using a common reverse primer and sequential HIV-specific forward primers.

The final amplicon was digested with EcoRI-HF, purified, and self-ligated with T4 ligase for inverse PCR. Ligation efficiency was assessed by qPCR. A final PCR step added Illumina adapters Libraries were gel-verified, purified, and prepared for paired-end sequencing using the NEBNext Ultra II DNA Library Prep Kit. Indexed libraries were pooled and sequenced on an Illumina NovaSeq 6000 SP system (PE150), generating ∼10 million reads per sample.

### DNA barcode and integration site data analysis

Barcode and UMI sequences were extracted using their flanking regions, and integration sites were defined as the sequence between the HIV-1 LTR and the L-shaped adapter as previously described^33^. Reads mapping to the NFNSX plasmid were excluded as contamination, and reads <10 nt were classified as linear unintegrated products. Remaining sequences were aligned to hg38 (Ensembl 108) or NFNSX using Bowtie2 and annotated as integrated, auto-integrated, or circular (immediately downstream of the HIV-1 LTR). True UMIs were identified by separating the bimodal count distribution (normal for true UMIs, Poisson for sequencing errors). For each true UMI, the most frequent barcode and integration site were assigned, yielding one provirus molecule per UMI.

### Statistical analysis

Data is presented using Prism 10 software (Graphpad). *P* < 0.05 was considered significant. Statistical details can be found in the corresponding figure legends. Flow cytometry analyses were performed using FlowJo 10.6.0. Deep sequencing analysis was performed using Bowtie2^45^, Jupyter, Bedtools, deepTools, SciPy, and BioPython.

## Acknowledgements

We acknowledge support from the National Institutes of Health (AI155232 to J.T.K. and R01AI161803 to C.A.B. and J.A.Z.), the California HIV/AIDS Research Program (H24BD7864 to J.T.K.), National Center for Advancing Translational Sciences UCLA CTSI Grant (UL1TR001881 to J.T.K.), and the UCLA-CDU Center for AIDS Research (AI152501). This research was supported in part by the CRISPR for Cure Martin Delaney Collaboratory for HIV cure UM1 AI164568-01 and cofunded by NIAID, NIMH, NIDA, NINDS, NIDDK, and NHLBI. The UCLA AIDS Institute, the McCarthy Family Foundation, and the James B. Pendleton Charitable Trust also provided support. ART was generously provided by Merck (RAL) and Gilead Sciences (TDF and FTC). Funding agencies did not play a specific role in design, analysis, decision to publish, or preparation of this manuscript.

## Contributions

Data acquisition, Y.S., I.A., W.H., M.K., A.G., H.C., M.D., J.T.K.; experimental design, Y.S., J.T.K.; data interpretation and analysis, Y.S., I.A., J.A.Z., J.T.K; preparation and writing of manuscript, Y.S., I.A., C.S., C.A.B., J.A.Z., and J.T.K.

## Data availability

All data supporting the findings of this study are available within the paper and its Supplementary Information. Source data are provided with this paper.

## Code availability

All original codes have been downloaded onto github at https://github.com/yuanshi-ys/D1D2NK_HIV.

