## Supplemental Figures for "NK Cells Engineered with a Chimeric Antigen Receptor Delay HIV Rebound and Reshape HIV Reservoir Composition"

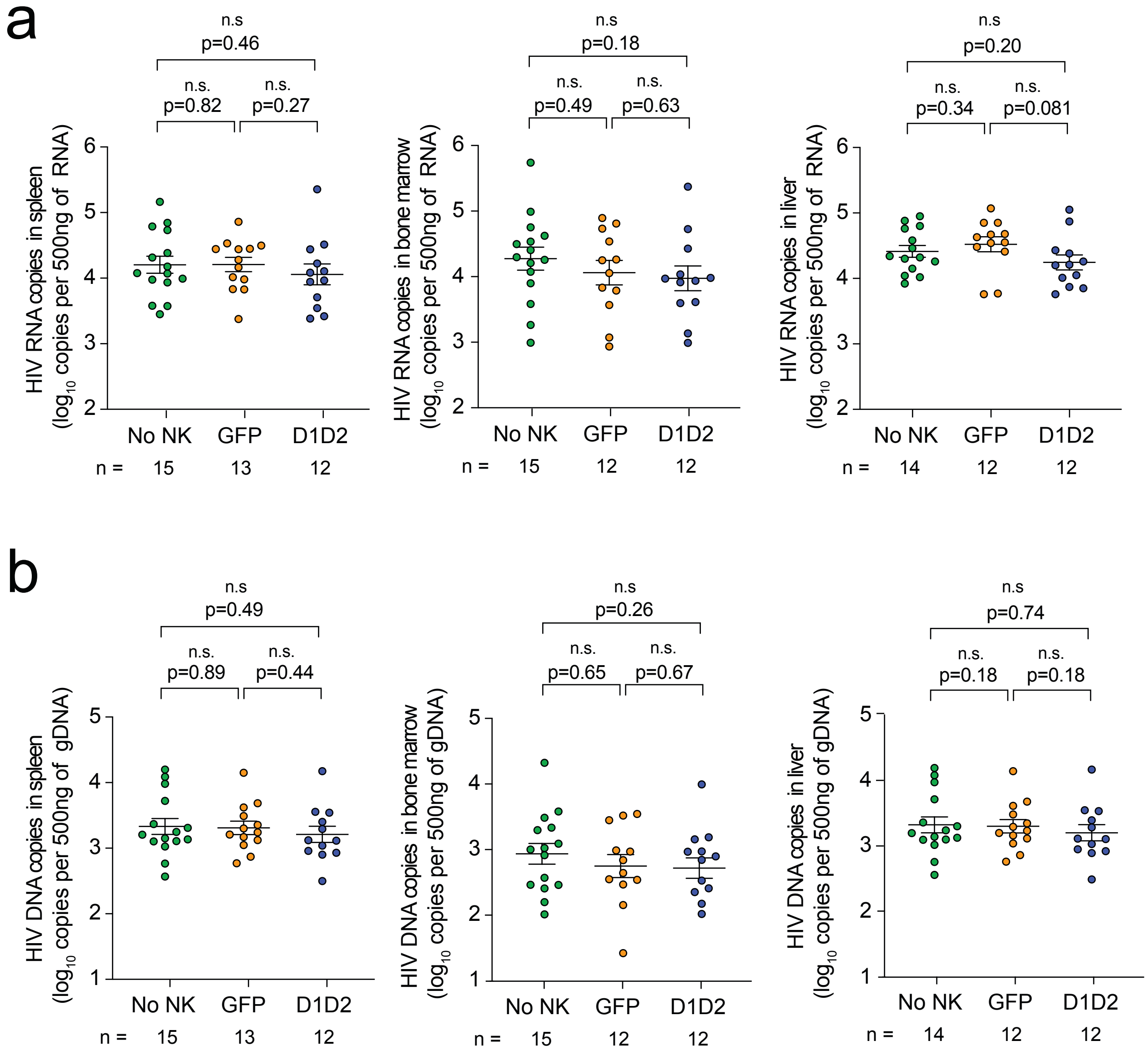


**Figure S1. Viral RNA and DNA levels during rebound infection at necropsy. a**, **b**) Cell-associated HIV RNA (**a**) and DNA (**b**) by RT-PCR and qPCR, respectively, from the spleen, bone marrow, and liver of mice during rebound infection at necropsy. n represents the number of mice. Shown are mean ± SEM. P values were determined by Mann-Whitney test.

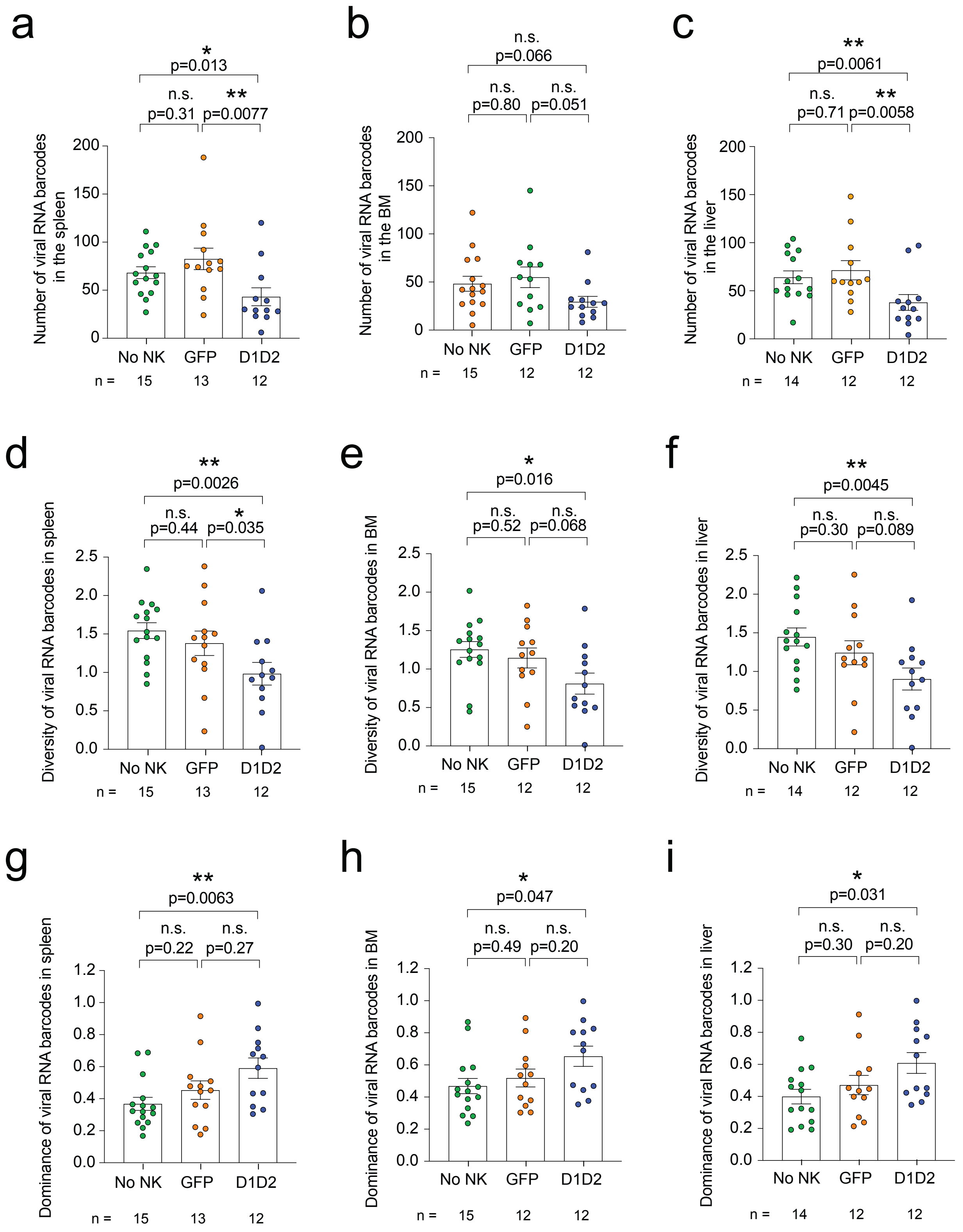


**Figure S2. Quantification of rebounding viral RNA barcodes, genetic diversity, and clonal dominance in spleen, bone marrow, and liver. a-c**) Number of rebounding viral RNA barcodes quantified by deep sequencing in the spleen (**a**), bone marrow (**b**), and liver (**c**). **d-f**) Genetic diversity of rebounding viral RNA barcoded measured using Shannon's diversity in the spleen (**d**), bone marrow (**e**), and liver (**f**). **g-i**) Dominance of viral RNA barcodes measured using Simpson's index in the spleen (**g**), bone marrow (**h**), and liver (**i**). n represents the number of mice. Shown are mean ± SEM. P values by two-tailed Mann-Whitney test.

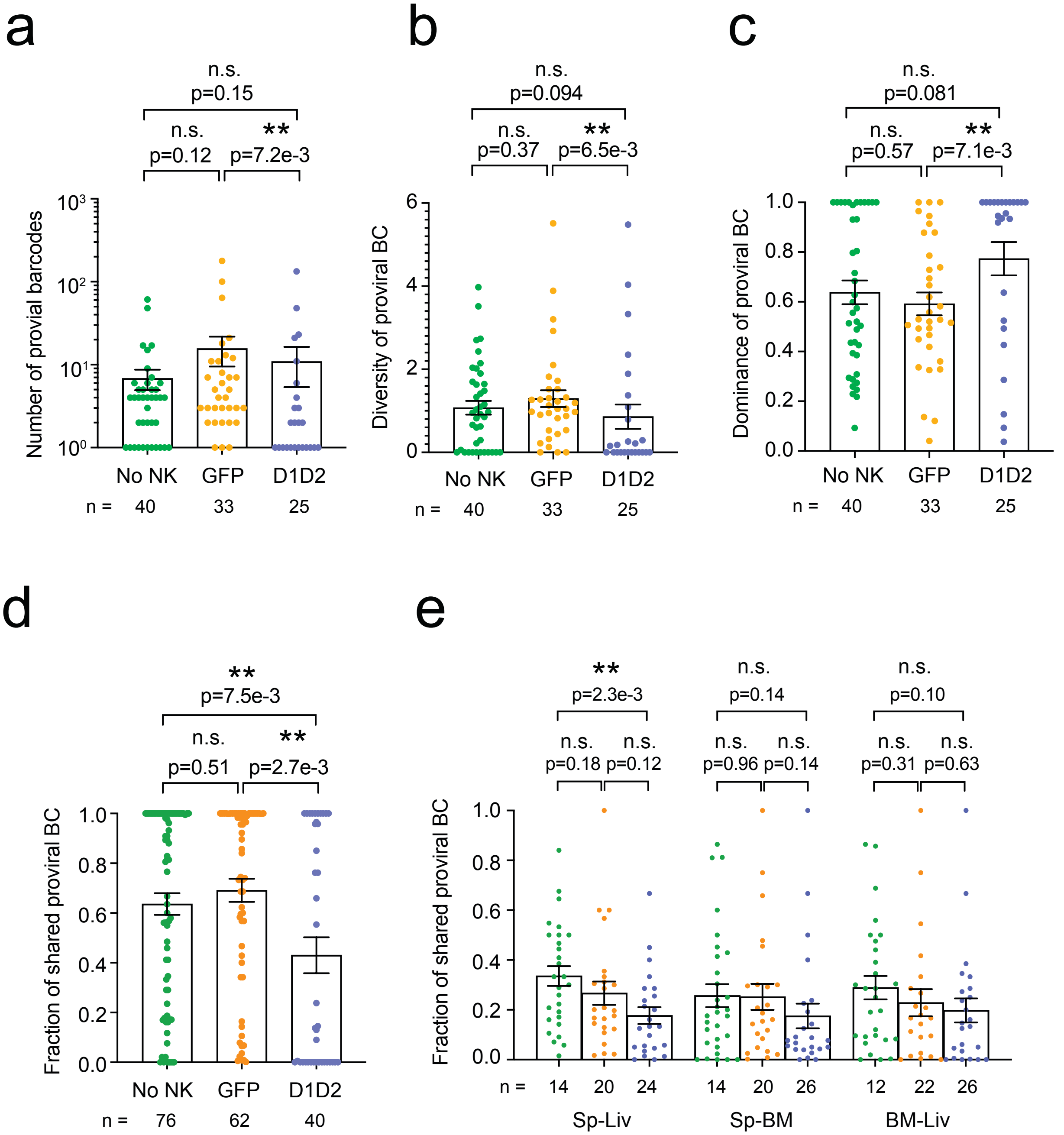


**Figure S3. Proviral barcode metrics and inter-organ clonal dynamics. a-c**) The number (**a**), genetic diversity (**b**) and dominance (**c**) of all proviral barcodes were quantified by deep sequencing. **d**) The fraction of all shared proviral barcodes between organ pairs were quantified by overlap index. **e**) The fraction of shared proviral barcodes associated with rebound viremia by organ pair. n represents the number of organs (**a-c**). n represents the number of organ pairs (**d, e**). Shown are mean ± SEM. P values by two-tailed Mann-Whitney test.

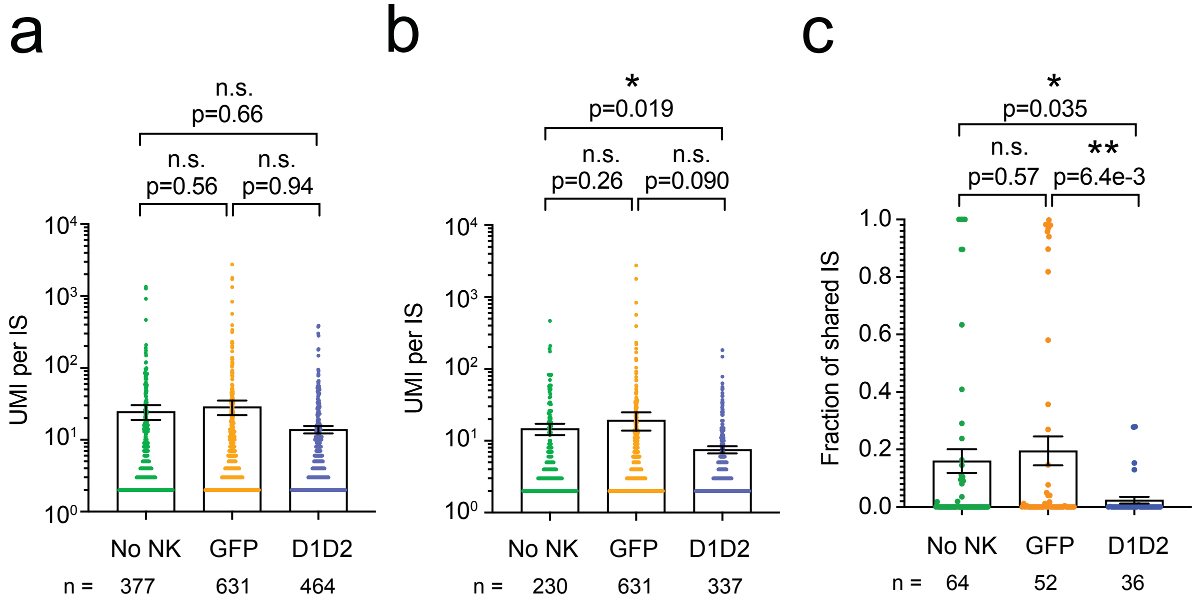


**Figure S4. Expanded cell clones and inter-organ sharing of integration sites. a**, **b**) The size of expanded cell clones harboring any provirus (**a**) or those with a provirus associated with rebound viremia (**b**). **c**) The fraction of shared integration sites among expanded clones between organ pairs was quantified by overlap index. n represents the number of integration sites (**a**, **b**). n represents the number of organ pairs (**c**). Shown are mean ± SEM. P values by two-tailed Mann-Whitney test.

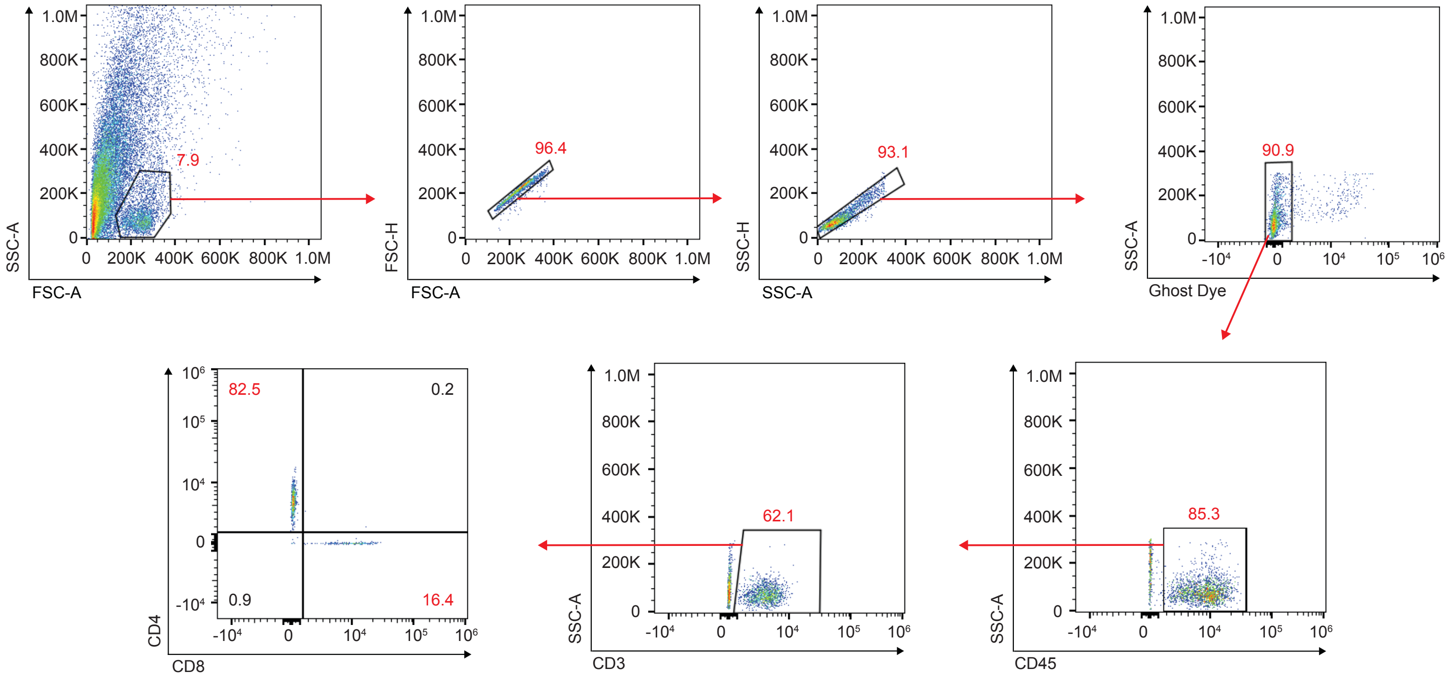


**Figure S5. Flow cytometry gating.** Representative flow cytometric analysis of human immune cells ~10 weeks post-CD34+ HSC injection into NSG mice.
